# Modulation of WNT, Activin/Nodal and MAPK Signaling Pathways Increases Arterial Hemogenic Endothelium and Hematopoietic Stem/Progenitor Cell Formation During Human iPSC Differentiation

**DOI:** 10.1101/2023.02.21.529379

**Authors:** Yongqin Li, Jianyi Ding, Daisuke Araki, Jizhong Zou, Andre Larochelle

## Abstract

Several differentiation protocols enable the emergence of hematopoietic stem and progenitor cells (HSPCs) from human induced pluripotent stem cells (iPSCs), yet optimized schemes to promote the development of HSPCs with self-renewal, multilineage differentiation and engraftment potential are lacking. To improve human iPSC differentiation methods, we modulated WNT, Activin/Nodal and MAPK signaling pathways by stage-specific addition of small molecule regulators CHIR99021, SB431542 and LY294002, respectively, and measured the impact on hematoendothelial formation in culture. Manipulation of these pathways provided a synergy sufficient to enhance formation of arterial hemogenic endothelium (HE) relative to control culture conditions. Importantly, this approach significantly increased production of human HSPCs with self-renewal and multilineage differentiation properties, as well as phenotypic and molecular evidence of progressive maturation in culture. Together, these findings provide a stepwise improvement in human iPSC differentiation protocols and offer a framework for manipulating intrinsic cellular cues to enable *de novo* generation of human HSPCs with functionality *in vivo*.

**Significance Statement:** The ability to produce functional HSPCs by differentiation of human iPSCs *ex vivo* holds enormous potential for cellular therapy of human blood disorders. However, obstacles still thwart translation of this approach to the clinic. In keeping with the prevailing arterial-specification model, we demonstrate that concurrent modulation of WNT, Activin/Nodal and MAPK signaling pathways by stage-specific addition of small molecules during human iPSC differentiation provides a synergy sufficient to promote arterialization of HE and production of HSPCs with features of definitive hematopoiesis. This simple differentiation scheme provides a unique tool for disease modeling, in vitro drug screening and eventual cell therapies.

## Introduction

Hematopoietic stem and progenitor cell (HSPC) transplantation is the most established cellular replacement therapy for a number of hematological diseases. However, the scarcity of human leukocyte antigen (HLA)-matched donors and insufficient numbers of long-term repopulating hematopoietic stem cells (HSCs) in donor umbilical cord blood (UCB) units often limit the ability to carry-out allogeneic HSPC transplant procedures. In the context of autologous gene therapy applications, sufficient functional HSCs from bone marrow (BM) or mobilized peripheral blood (MPB) cell collections are also commonly unavailable in patients with impaired hematopoiesis, such as BM failure syndromes and chronic inflammatory disorders [1–3]. Current protocols to augment cell dose by expansion of human HSCs *ex vivo* are inefficient [4–6]. *De novo* generation of human HSPCs is a tractable alternative for the development of cell-based therapies when primary HSPCs are limited. One approach relies on reprogramming adult somatic cells to an induced pluripotent stem cell (iPSC) state by enforced expression of a defined set of transcription factors [7–9]. For autologous HSPC-based gene therapies, programmable CRISPR/Cas9 nuclease systems may be used for targeted integration of a wild-type therapeutic open reading frame construct within patients’ iPSCs. Although efficiencies of genetic correction are generally low, disease-free iPSC clones can be selected and readily expanded in culture. Induced pluripotent stem cells are then subjected to differentiation into a theoretically infinite supply of HSPCs for transplantation [10,11].

E*x vivo* iPSC differentiation approaches generally aim at recapitulating the natural hematopoietic developmental process that occurs during ontogeny. Induction of the mesoderm primordial germ layer marks the onset of hematopoiesis in the embryo. Mesodermal progenitors then specify to hematoendothelial fates and establish a series of temporally and spatially distinct blood systems or “waves” to support the growing embryo. The first wave of hematopoietic potential, designated “primitive”, is highly restricted and primarily results in the emergence of erythroid, macrophage and megakaryocyte progenitors. A second wave of hematopoiesis, dubbed “definitive”, supplies lymphoid and erythro-myeloid progenitor cells. The third wave of blood formation is also viewed as “definitive” but is uniquely marked by the appearance of multilineage engrafting HSCs primarily within the ventral wall of the dorsal aorta in the aorto-gonad-mesonephros (AGM) region of the embryo proper [12–15]. Within the AGM, HSCs arise from hemogenic endothelium (HE), a unique subset of unipotent vascular endothelium fated to a blood lineage identity [16–18]. The mechanism by which HE cells gain hematopoietic potential and morphology at the expense of their endothelial identity is known as endothelial-to-hematopoietic transition (EHT) [19–22]. During development, definitive HSCs were shown to arise from HE lining arteries, but not veins, evoking the possibility that arterial specification of HE is necessary to initiate the definitive hematopoietic program within the embryo [23–27]. Following their emergence in the AGM, nascent HSCs undergo maturation through complex interactions with niche constituents of various hematopoietic sites during development [28]. Hence, successful iPSC differentiation protocols are contingent on faithfully reproducing the early stages of mesodermal induction and specification, and the definitive third wave of hematopoiesis during which HE cells emerge, undergo arterial fate specification and progress through an EHT phase for the generation of mature HSCs with functionality *in vivo*.

Seminal studies have uniformly shown that stepwise addition of cytokines and morphogens alone during differentiation induces pluripotent stem cells (PSCs) to develop a yolk sac-like primitive hematopoietic program, with limited arterial HE activation and inefficient production of HSCs with long-term engraftment capability [29–37]. Early mesodermal stage activation of WNT cellular signaling and repression of the Activin/Nodal pathway by one-time addition of CHIR99021 (CHIR) and SB431542 (SB) small molecules, respectively, are now customarily used to block primitive hematopoiesis and enhance the definitive hematopoietic program in human PSC differentiation protocols [38,39]. Moreover, formation of HE expressing arterial markers (e.g., DLL4 or CXCR4) can be augmented by supplementing culture medium at the mesodermal phase of human PSC differentiation with Ly294002 (LY), a small molecule inhibitor of PI3K kinase, for indirect activation of MAPK cellular signaling [23]. The impact of MAPK signaling activation on arterial HE specification is primarily mediated through stimulation of the NOTCH pathway, a critical determinant of arterial specification in embryonic vasculature [40–42]. Only arterialized HE populations produce a definitive-type hematopoiesis in culture, but this approach alone is insufficient to promote formation of engraftable HSCs *in vitro*.

In this study, we assessed whether a combined approach to modulate WNT, Activin/Nodal and MAPK cellular signaling might provide a synergy sufficient to further promote definitive hematopoiesis and the generation of engrafting HSCs. We found that manipulation of these pathways by addition of CHIR, SB and LY during human iPSC hematopoietic differentiation significantly enhanced formation of arterial HE and derivation of human HSPCs with self-renewal properties, multilineage differentiation capacity and evidence of progressive maturation in culture.

## Materials and Methods

### Generation and culture of human iPSCs

Human peripheral blood CD34^+^ cells were G-CSF mobilized and apheresed from three independent healthy donors after informed consent, in accordance with the Declaration of Helsinki, under an Institutional Review Board-approved clinical protocol (NCT00027274). Human CD34^+^ cells from each donor were sorted to enrich for the most primitive HSC population (CD34^+^CD38^−^ cells) using BD FACSAria™ II or BD FACSAria™ Fusion instruments with a 100 μm nozzle. Sorted CD34^+^CD38^−^ cells were reprogrammed into iPSCs (MCND-TEN-S2, HT914 and HT915 iPSC lines) using the integration-free CytoTune 2.0 Sendai virus reprogramming kit (A16517, Thermo Fisher Scientific) as previously reported [43–45]. Data reported in this study were obtained with MCND-TEN-S2 (registered at https://hpscreg.eu/cell-line/RTIBDi001-A) and reproducibility of our results was confirmed with HT914 and HT915 where indicated. Pluripotency was confirmed by teratoma formation assay and flow cytometry for TRA-1–60 and NANOG markers as previously described [46,47]. Chromosomal integrity was verified by karyotyping using GTW-banded chromosome method at the WiCell Research Institute (Madison, WI, USA). Twenty metaphase cells were analyzed for each iPSC clone. iPSCs were maintained in tissue culture plates coated with Matrigel® (#354230, Corning) in Essential 8™ (E8) Medium (#A1517001, Thermo Fisher). Culture medium was changed daily and iPSCs were split every three to four days with 0.5mM EDTA in phosphate buffered saline (PBS) at various ratios.

### Hematopoietic differentiation of human iPSCs

Human iPSCs were differentiated for 12 days using the STEMdiff™ Hematopoietic Kit (#05310, STEMCELL Technologies, Inc.) [43]. Briefly, one day before differentiation, iPSCs were split into small clusters using PBS/EDTA, and cluster concentrations were calculated. A total of 20–35 clusters were transferred into each well of a Matrigel-coated 12-well plate and cultured overnight in E8 medium with 1.25 μM ROCK inhibitor (#Y0503, Sigma). On day 0 of differentiation, medium A (containing basic fibroblast growth factor [bFGF], bone morphogenetic protein 4 [BMP4] and vascular endothelial growth factor A [VEGFA]) was added to promote mesodermal differentiation and a half medium A change was done on day 2. On day 3 of differentiation, supernatant was removed and hematopoietic differentiation medium B (containing bFGF, BMP4, VEGFA, stem cell factor [SCF], FMS-line tyrosine kinase 3 ligand [Flt3L] and thrombopoietin [TPO]) was added, followed by half-medium changes on days 5, 7, 9 and 11. In select experiments, 3 μM CHIR99021 (#SML1046, Sigma) and 6 μM SB431542 (#S4317, Sigma) were added on day 2 for 24–36 h, and 3 μM LY294002 (#70920, Cayman Chemicals) was added from day 3 to day 6 of differentiation unless otherwise indicated.

### Collection of adherent and non-adherent cells differentiated from iPSCs

On days when flow cytometry analysis was performed, both CD43^+^CD45^+/−^ hematopoietic (suspension) and CD43^−^CD45^−^ non-hematopoietic (adherent monolayer) fractions were harvested and combined for analysis. Hematopoietic cells were collected first by vigorous pipetting. The remaining non-hematopoietic population was washed once with PBS, incubated with Accutase™ (#07920, STEMCELL Technologies, Inc.) for 10 minutes at 37°C, and vigorously pipetted up and down to ensure complete recovery with PBS/10% fetal bovine serum (FBS). Cells were then combined and filtered through a 40 μm cell strainer, counted, and prepared for analysis.

### Flow cytometry and fluorescence-activated cell sorting (FACS)

Cells were stained with antibodies (Table S1) following manufacturer’s instructions, and analyzed on an LSRII Fortessa analyzer (Becton Dickinson, BD). All flow data were analyzed using FlowJo 10.7 Software. For gene expression and colony forming unit (CFU) assays, cell populations were sorted on BD FACSAria™ II or BD FACSAria™ Fusion instrument with a 100 μm nozzle.

### CFU assay

Day 12 suspension hematopoietic cells were collected, sorted and resuspended in Iscove’s Modified Dulbecco’s Medium (IMDM, Sigma Aldrich) supplemented with 2% FBS. Human CFU assays were performed as per manufacturer’s instructions (#04445, STEMCELL Technologies, Inc.). Briefly, 9000 CD43^+^CD45^+/−^CD34^+^ cells (DMSO and LY groups) or 4500 cells (CHIR/SB and CHIR/SB/LY groups) were suspended in 300 uL IMDM/2% FBS, which was then added to 3 mL methylcellulose and vortexed. A volume of 1.1 mL was plated onto 35 mm tissue-culture dishes (#353001, Corning) for a total of 1500 to 3000 cells per plate. Colonies were scored manually 12-14 days following plating. In select experiments, colony-forming activity of iPSC-derived HSPCs was assessed after a 10-day culture period (37°C, 21% O_2_, 5% CO_2_) at a cell concentration of 5 × 10^5^/mL in StemSpan Serum-Free Expansion Medium II (STEMCELL Technologies) supplemented with 100 ng/mL of human recombinant SCF, Flt3L, and TPO (PeproTech).

### CFU replating assay

For colony replating experiments, 2 weeks after the primary plating, colonies from two plates were pooled, washed with PBS, and resuspended in 300 uL IMDM/2% FBS. Cells were then plated in new methylcellulose medium at a concentration of 25,000 cells per mL onto 35 mm tissue-culture dishes. Colonies were scored manually 14 days following secondary plating.

### Gene expression by real-time qPCR

Total RNA was extracted from sorted cells using RNeasy Micro Kit (#74004, Qiagen). Complementary DNA was synthesized with PrimeScript™ RT Reagent Kit (#RR037A, Takara) at 37°C for 15 min and de-activated at 85°C for 5 sec. PCR with reverse transcription was performed using PowerUp™ SYBR™ Green Master Mix (#A25777, Thermo Fisher Scientific) on a BioRad C1000 touch system. The fold difference in mRNA expression between treatment groups was determined using the ΔΔCt method. All samples were multiplexed to include an internal GAPDH control. Primers and probes used for real-time qPCR are listed in Table S2.

### Mouse transplantation

6- to 12-week-old NOD.Cg-Kit^W-41J^ Tyr^+^ Prkdc^scid^ Il2rg^tm1Wjl^/ThomJ mice (NBSGW, stock #026622) were purchased from Jackson Laboratory. Animals were housed and handled in accordance with the guidelines set by the Committee on Care and Use of Laboratory Animals of the Institute of Laboratory Animal Resources, National Research Council (DHHS publication No. NIH 85–23), and the protocol was approved by the Animal Care and Use Committee of the NHLBI. Day 12 CD34^+^ hematopoietic cells were purified via magnetic-activated cell sorting (MACS) using human CD34 microbeads (#130-046-702, Miltenyi Biotec). As previously reported [48], 48 hours prior to the infusion of human cells, mice received 10 mg/kg Busulfex (PDL BioPharma) for conditioning by intraperitoneal injection. For homing studies, 2×10^6^ sorted CD34^+^ hematopoietic cells were resuspended in PBS and injected by intravenous injection into the tail vein of NBSGW mice. Homing of infused cells within the bone marrow and spleen was evaluated 17-24 hours after injection. For long-term transplantation studies, sorted CD34^+^ hematopoietic cells were resuspended at 2×10^6^ cells per 25 μL medium and transplanted by intra-femoral injection. Before transplantation, mice were temporarily anesthetized with isoflurane inhalation. A 26G needle was used to create a tunnel into the bone marrow cavity of the femur. Cells were transplanted into the tunnel using a 28.5G insulin needle. Up to 100 μL peripheral blood was collected every 4 weeks through 16 weeks. Bone marrow was collected 16 weeks post-transplantation and stained with human CD45-PE antibodies (Table S1) to quantify human cell engraftment. Investigators were blinded for the analysis of mice.

### Statistical analysis

Results were analyzed with GraphPad Prism Software (version 9.0.2), using two-tailed unpaired t-test (panels B, D, E and F) and one-way ordinary ANOVA test with Dunnett correction. Results are displayed as mean ± standard error of the mean (SEM). ns, not significant, * p<0.05, ** p<0.01, *** p<0.001, and **** p<0.0001.

## Results

### Modulation of WNT, Activin/Nodal and MAPK signaling pathways enhances formation of arterial HE during human iPSC differentiation

To determine the impact of modulating WNT, Activin/Nodal and MAPK intracellular signaling on hematoendothelial formation in culture, human iPSCs reprogrammed from CD34^+^CD38^−^ cells of healthy volunteers were subjected to hematopoietic differentiation for 12 days using our previously reported monolayer culture system [43] with or without stage-specific addition of small molecule regulators of these pathways. In this platform, a non-hematopoietic CD43^−^CD45^−^ adherent monolayer rapidly forms under conditions that support mesodermal induction (day 0 to 3). With the subsequent addition of hematopoietic cytokines that promote mesodermal specification to hematoendothelial fates and HSPC formation/maturation (day 3 to 12), HE progenitors develop (peak HE production at day 6 of culture) and CD43^+^CD45^+/−^ hematopoietic cells arise from the monolayer before their eventual release within the supernatant fraction (peak HSPC production at day 12 of culture) [43] (Fig. 1A).

**Figure 1.**
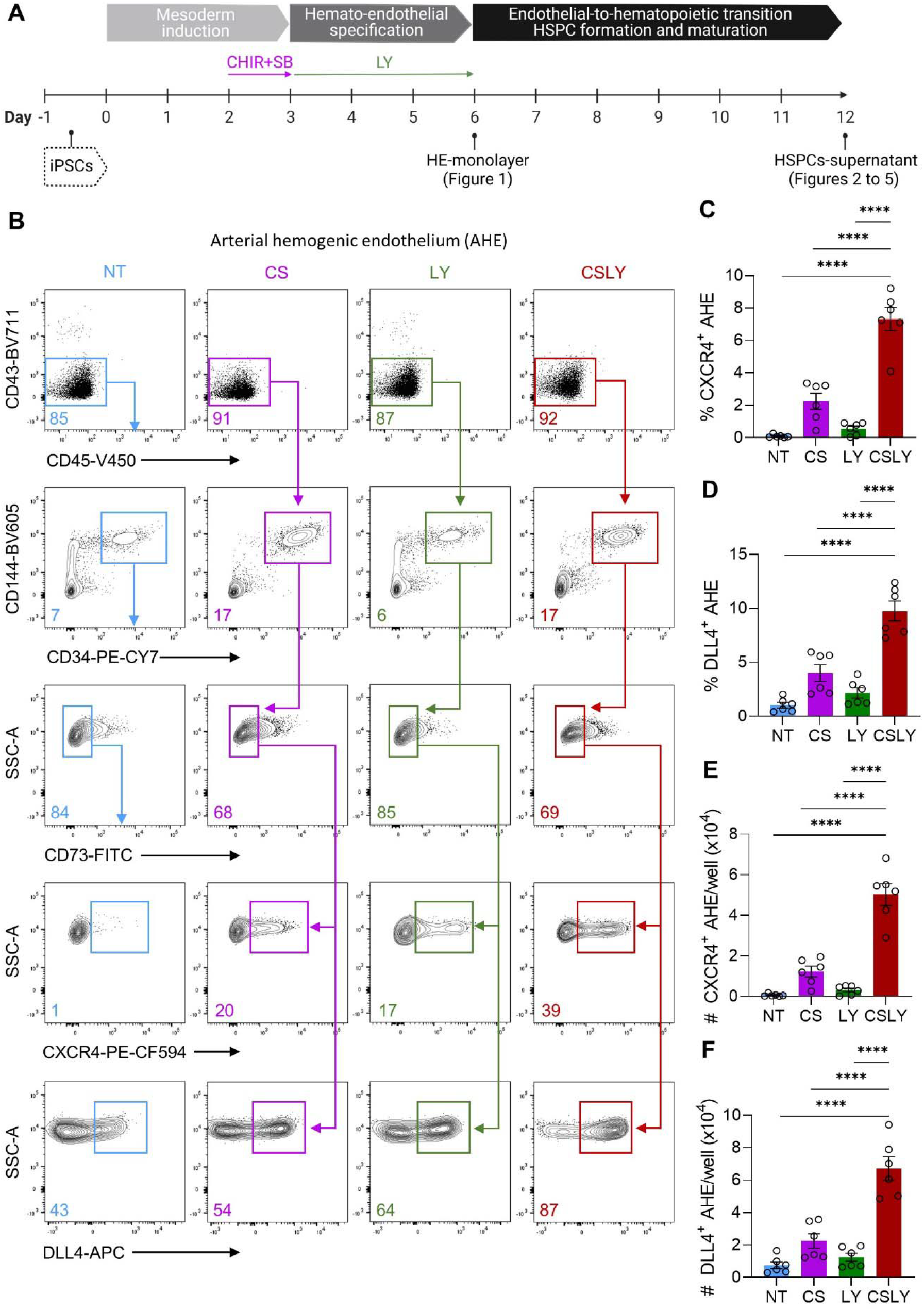
Modulation of WNT, Activin/Nodal, and MAPK signaling pathways during human iPSC differentiation enhances formation of arterial hemogenic endothelium. **(A)** Schematic of the experimental design for hematopoietic differentiation of human iPSCs, including mesodermal induction (day 0 to 3), hemato-endothelial specification (day 3 to 6) and hematopoietic differentiation (day 6 to 21). Maximal production of hemogenic endothelium (HE) within the CD43^−^CD45^−^ adherent non-hematopoietic monolayer was observed at day 6 of culture, and peak formation of hematopoietic stem and progenitor cells (HSPCs) was detected within the CD43^+^CD45^+/−^ supernatant hematopoietic fraction at day 12 of differentiation. Unless otherwise indicated, WNT agonist CHIR99021 (CHIR) and Activin/Nodal antagonist SB431542 (SB) were supplemented on day 2 of differentiation for 24 hours, and MAPK agonist LY294002 (LY) was added from day 3 to 6 of culture. All analyses were performed with cells collected at day 6 of differentiation in the absence of small molecule adjunct treatment (non-treated, NT) or in the presence of CHIR/SB (CS), LY or CHIR/SB/LY (CSLY). **(B)** Representative flow cytometry plots of arterial HE (AHE)-defining markers (CD43^−^CD45^−^CD144^+^CD34^+^CD73^−^ CXCR4^+^ or DLL4^+^). **(C)** Percentages of CXCR4^+^ AHE cells. **(D)** Percentages of DLL4^+^ AHE cells. (**E)** Absolute numbers of CXCR4^+^ AHE cells per culture well. **(F)** Absolute numbers of DLL4^+^ AHE cells per culture well. Data shown in this figure were obtained with human iPSC line MCND-TEN-S2. Reproducibility of data was confirmed with human iPSC lines HT914 (Fig. S1). In panels C-F, data are displayed as mean ± standard error of the mean (SEM) of 6 independent experiments. One-way ordinary ANOVA test with Dunnett correction was used. **** p ≤ 0.0001. Associated with Figs. S1 and S2.

As initially demonstrated in other PSC culture protocols [38,39], we previously confirmed that supplementing CHIR (WNT agonist) and SB (Activin/Nodal repressor) at day 2 of iPSC differentiation for a period of 24 hours suppresses primitive hematopoiesis and promotes definitive HSPC formation in our culture system [43]. To assess whether LY-mediated activation of MAPK signaling at the mesodermal stage of development also enhances HE arterial specification in our differentiation approach, we supplemented LY from day 3 through the end of mesodermal induction/specification (day 6). Production of arterial HE was measured by interrogating the CD43^−^CD45^−^ non-hematopoietic cell fraction for HE-defining markers, including CD144^+^ (VE-cadherin^+^), CD34^+^ and lack of CD73 expression (CD73^−^), as well as markers previously associated with arterialization of HE (CXCR4^+^ and DLL4^+^) [23,25,49]. Compared to control cultures containing no CHIR/SB/LY (non-treated, NT) or supplemented with CHIR/SB or LY only, combined addition of CHIR, SB and LY led to a marked increase in percentages (Fig. 1B-D and Fig. S1) and numbers (Fig. 1E, F and Fig. S1) of CXCR4^+^ or DLL4^+^ arterial HE (CD43^−^ CD45^−^CD144^+^CD34^+^CD73^−^CXCR4^+^/DLL4^+^). The peak effect was observed on the 6^th^ day of differentiation and when LY was supplemented from day 3 through day 6 of culture (Fig. S2).

### Modulation of WNT, Activin/Nodal and MAPK signaling pathways promotes definitive HSPC formation during human iPSC differentiation

We next investigated whether the early increase in arterial HE formation observed in the presence of CHIR/SB/LY influenced hematopoietic development (Fig. 2A). Addition of CHIR/SB during differentiation decreased overall CD43^+^CD45^+/−^ hematopoietic cell numbers compared to untreated groups, but supplementation of culture medium with LY in combination with CHIR/SB offset this effect and LY alone had no impact on total cell numbers in culture (Fig. 2B). Moreover, a rise in percentages (Fig. 2C) and numbers (Fig. 2D) of CD34^+^ progenitors was observed within the hematopoietic population at day 12 of differentiation with CHIR/SB/LY compared to controls. In CFU assays, the frequency of progenitors with multilineage differentiation capacity was similar between control groups but significantly increased in the presence of CHIR/SB/LY (Fig. 2E, F).

**Figure 2.**
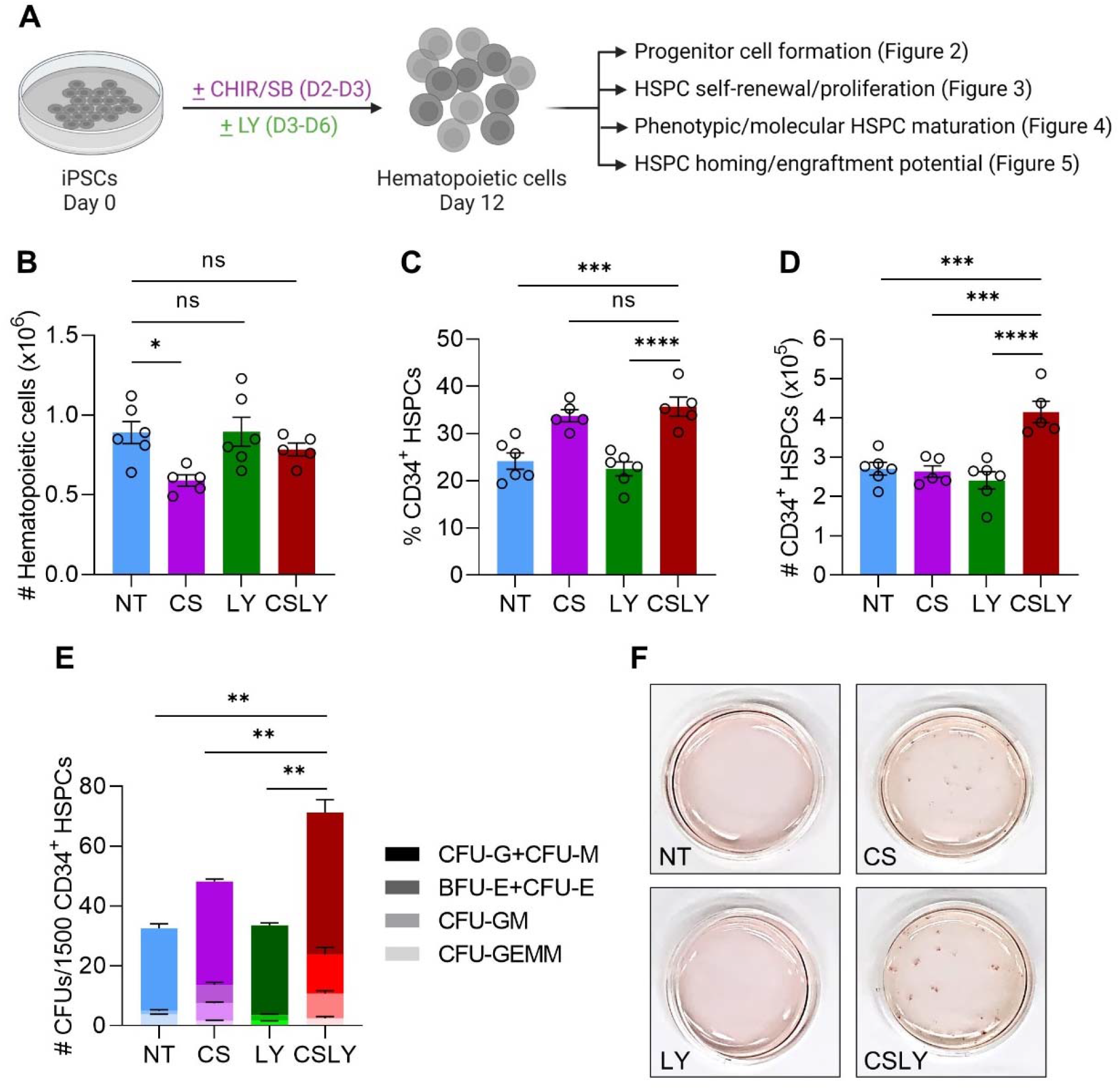
Modulation of WNT, Activin/Nodal, and MAPK signaling pathways during human iPSC differentiation enhances formation of hematopoietic progenitors. **(A)** Experimental scheme. Human iPSCs were differentiated as illustrated in Fig. 1A in the absence of small molecule adjunct treatment (non-treated, NT) or in the presence of CHIR/SB (CS), LY or CHIR/SB/LY (CSLY). Hematopoietic progenitor activity of cells released within the culture supernatant at day 12 of differentiation with indicated treatments was evaluated by flow cytometry and colony forming unit (CFU) assay. (**B)** Absolute numbers of CD43^+^CD45^+/−^ hematopoietic cells per culture well. **(C)** Percentages of CD34^+^ HSPCs. **(D)** Absolute numbers of CD34^+^ HSPCs per culture well. **(E)** Numbers of myeloid (CFU-G, CFU-M, CFU-GM and CFU-GEMM) and erythroid (CFU-E, BFU-E) colonies per 1500 CD34+ cells purified by FACS. **(F)** Representative images of CFU plates presented in panel E demonstrating a significant increase in colony counts by addition of CSLY during iPSC differentiation. Data shown in this figure were obtained with human iPSC line MCND-TEN-S2. Reproducibility of data was confirmed with human iPSC lines HT914 and HT915 (Fig. S3). In panels B-E, data are displayed as mean ± SEM of 3 to 6 independent experiments. One-way ordinary ANOVA test with Dunnett correction was used. ns, not significant, * p ≤ 0.05, ** p ≤ 0.01, *** p ≤ 0.001, **** p ≤ 0.0001. Associated with Fig. S3.

Next, we assessed whether the CHIR/SB/LY combination enabled the emergence of third wave definitive HSPCs with self-renewal (Fig. 3), progressive maturation (Fig. 4) and homing/engraftment potential during iPSC differentiation (Fig. 5). To gain insights into the self-renewal and proliferative capacities of iPSC-derived HSPCs, hematopoietic colonies that formed in primary CFU cultures were pooled and replated in secondary clonogenic assays (Fig. 3A, top). Notably, we observed a significant (2.2- to 6.2-fold) increase in replating efficiency from CD34^+^ HSPCs arising from CHIR/SB/LY-supplemented cultures compared to control groups (Fig. 3B). To further corroborate these findings, day 12 CD34^+^ cells derived from each iPSC differentiation condition were cultured for 10 days in cytokine-supplemented medium before quantification of CFU activity in clonogenic progenitor assays (Fig. 3A, bottom). As previously shown, iPSC-differentiated HSPCs that lack self-renewal and proliferative abilities differentiate and lose their colony-forming potential in extended culture [50]. Using this experimental scheme, only rare colonies were detected from untreated or LY-supplemented iPSC differentiation conditions. HSPCs derived from CHIR/SB-containing iPSC cultures and differentiated for 10 days *in vitro* produced increased CFU numbers relative to untreated and LY groups, but the most substantial rise in colony-forming activity after a 10-day culture period was unequivocally observed with CD34^+^ HSPCs differentiated from iPSCs in the presence of CHIR/SB/LY (Fig. 3C).

**Figure 3.**
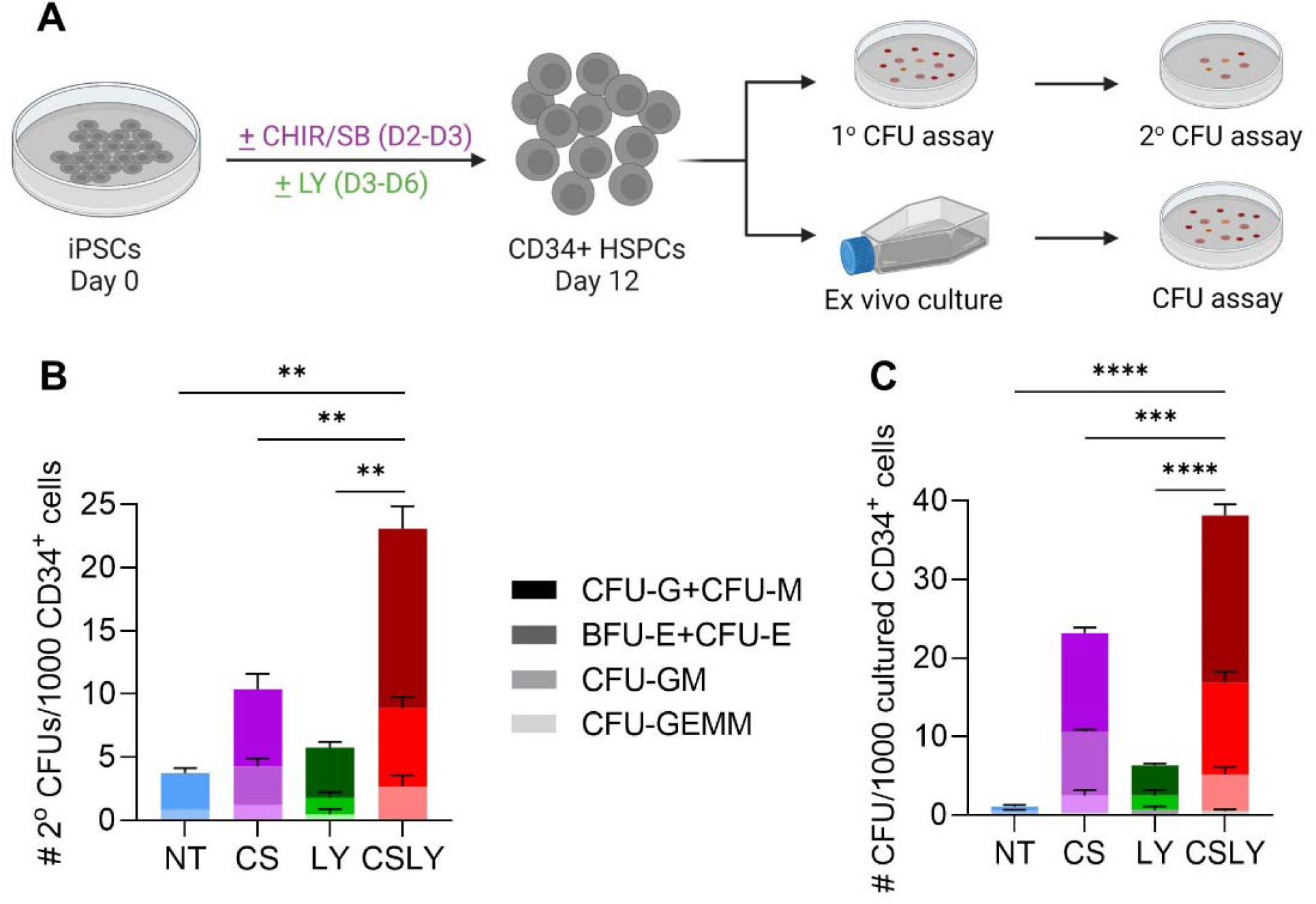
Modulation of WNT, Activin/Nodal, and MAPK signaling pathways during human iPSC differentiation enhances formation of HSPCs with self-renewal capacity. **(A)** Experimental scheme. Human iPSCs were differentiated as illustrated in Fig. 1A in the absence of small molecule adjunct treatment (non-treated, NT) or in the presence of CHIR/SB (CS), LY or CHIR/SB/LY (CSLY). Self-renewal capacity of HSPCs released within the culture supernatant at day 12 of differentiation with indicated treatments was evaluated by secondary clonogenic assay or CFU formation following extended (10-day) culture. (**B)** Secondary clonogenic assay. A total of 1000 CD34+ HSPCs purified by FACS were plated in primary CFU assays. After 12-14 days, colonies were scored, pooled and equal numbers of cells were replated for each condition. Secondary CFU plates were scored at day 12-14 for myeloid (CFU-G, CFU-M, CFU-GM and CFU-GEMM) and erythroid (CFU-E, BFU-E) colonies, and counts were normalized to the total number of cells from primary CFU plates. **(C)** CFU assay after extended culture. A total of 1000 CD34+ HSPCs purified by FACS were cultured for 10 days and subsequently plated in primary CFU assays. CFU plates were scored at day 12-14 for myeloid and erythroid colonies. Data shown in this figure were obtained with human iPSC line MCND-TEN-S2. Reproducibility of data was confirmed with human iPSC lines HT914 and HT915 (Fig. S4). In panels B and C, data are displayed as mean ± SEM of 3 independent experiments. One-way ordinary ANOVA test with Dunnett correction was used. ** p ≤ 0.01, *** p ≤ 0.001, **** p ≤ 0.0001. Associated with Fig. S4.

**Figure 4.**
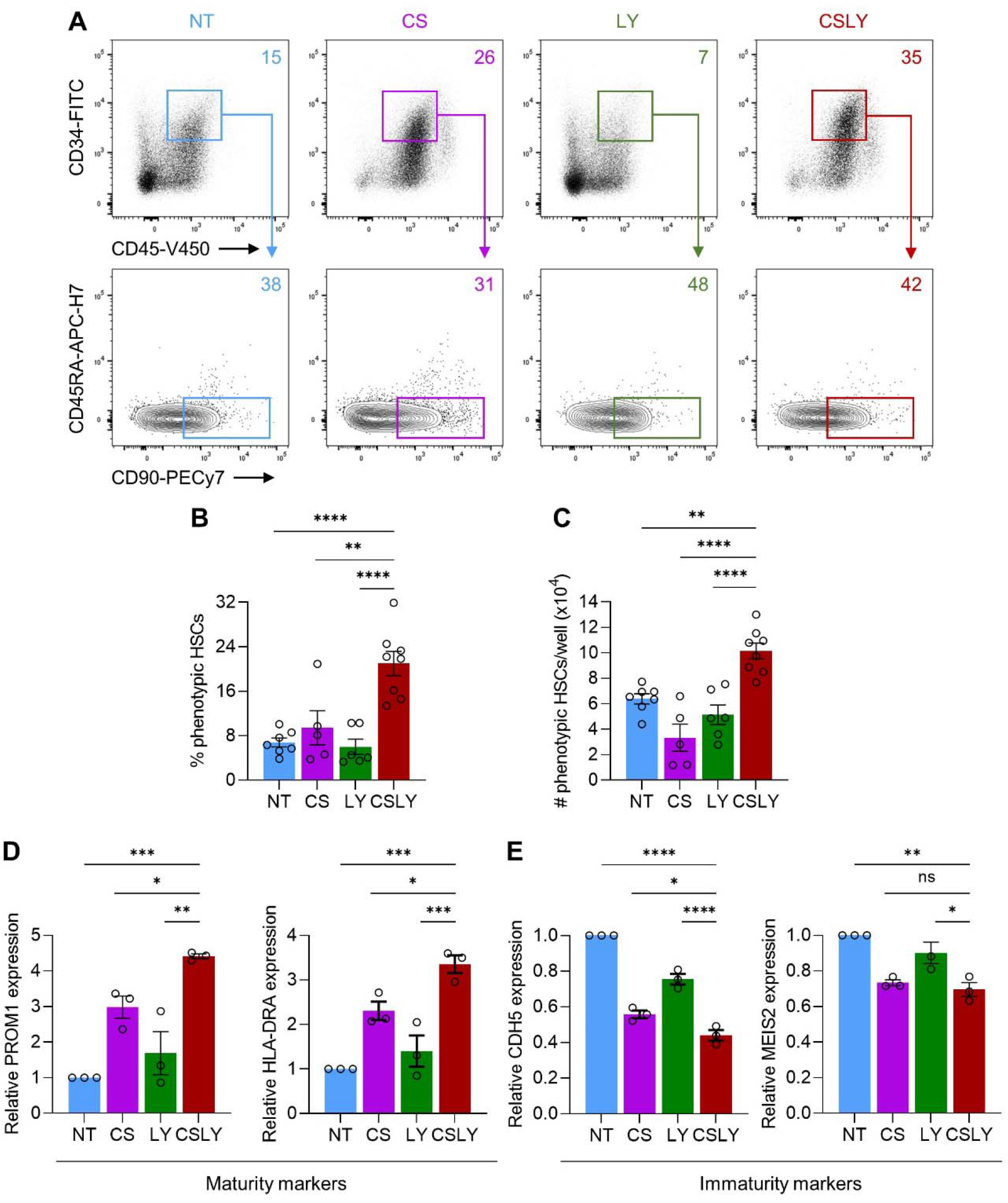
Modulation of WNT, Activin/Nodal, and MAPK signaling pathways during human iPSC differentiation enhances formation of HSPCs with phenotypic and molecular attributes of maturation. Human iPSCs were differentiated as illustrated in Fig. 1A in the absence of small molecule adjunct treatment (non-treated, NT) or in the presence of CHIR/SB (CS), LY or CHIR/SB/LY (CSLY). All analyses were performed with HSPCs collected at day 12 of differentiation with indicated treatments. **(A)** Representative flow cytometry plots of phenotypically defined HSCs (CD45^+^CD34^+^CD45RA^−^CD90^+^). **(B)** Percentages of phenotypically defined HSCs. **(C)** Absolute numbers of phenotypically defined HSCs per culture well. **(D)** Quantitative PCR analysis of PROM1 and HLA-DRA molecular markers associated with hematopoietic maturity in phenotypically defined HSCs. Data were normalized to the NT control group. **(E)** Quantitative PCR analysis of CDH5 and MEIS2 molecular markers associated with hematopoietic immaturity in phenotypically defined HSCs. Data were normalized to the NT control group. Data shown in this figure were obtained with human iPSC line MCND-TEN-S2. Reproducibility of data was confirmed with human iPSC line HT915 (Fig. S5). In panels B-E, data are displayed as mean ± SEM of 3 to 8 independent experiments. One-way ordinary ANOVA test with Dunnett correction was used. ns, not significant, * p ≤ 0.05, ** p ≤ 0.01, *** p ≤ 0.001, **** p ≤ 0.0001. Associated with Fig. S5.

**Figure 5.**
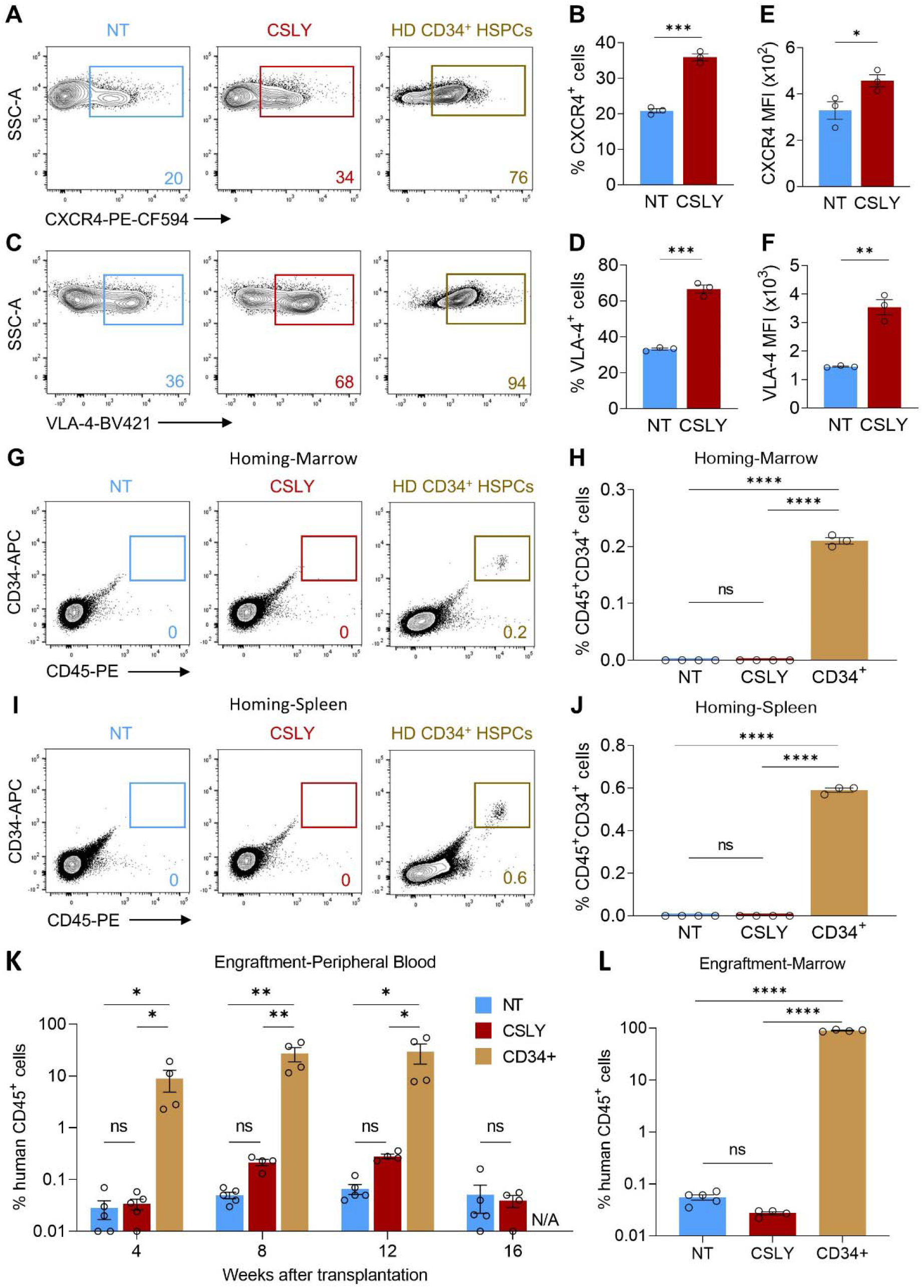
Modulation of WNT, Activin/Nodal, and MAPK signaling pathways during human iPSC differentiation is insufficient to enhance formation of HSPCs with functional properties *in vivo*. Human iPSCs were differentiated as illustrated in Fig. 1A in the absence of small molecule adjunct treatment (non-treated, NT) or in the presence of CHIR/SB/LY (CSLY). All analyses were performed using CD34^+^ HSPCs purified by FACS from the culture supernatant at day 12 of differentiation with indicated treatments or collected by G-CSF mobilization and apheresis of a healthy donor (HD). **(A-F)** Expression of CXCR4 and VLA-4 homing markers. Panels display representative flow cytometry plots of CXCR4 (A) or VLA-4 (C) expression, percentages of cells expressing CXCR4 (B) or VLA-4 (D), and mean fluorescence intensity (MFI) of CXCR4 (E) or VLA-4 (F) homing markers. **(G-J)** Early *in vivo* homing potential. Panels display representative flow cytometry plots and percentages of human CD45^+^CD34^+^ cells detected within the marrow (G, H) or spleen (I, J) of NBSGW mice at 24 hours post-injection. **(K-L)** Long-term *in vivo* engraftment potential. Panels display percentages of human CD45^+^ cells detected within the peripheral blood of NBSGW mice at the indicated times post-transplantation (K) or within the murine bone marrow at 4 months post-transplantation (L). Data shown in this figure were obtained with human iPSC line MCND-TEN-S2. In panels B, D, E and F, data are displayed as mean ± SEM of 3 independent experiments. In panels H, J, K and L, data are displayed as mean ± SEM of 3 to 5 mice for each group. Two-tailed unpaired t-test (panels B, D, E and F) and one-way ordinary ANOVA test with Dunnett correction (panels H, J, K and L) were used. ns, not significant, * p ≤ 0.05, ** p ≤ 0.01, *** p ≤ 0.001, **** p ≤ 0.0001. Associated with Fig. S6.

To determine the impact of CHIR/SB/LY on the extent of HSPC maturation at day 12 of differentiation, we first tracked the CD34^+^CD38^−^CD45RA^−^CD90^+^ cell surface phenotype customarily accepted to define HSPC populations highly enriched in HSCs with engraftment potential [51]. In iPSC cultures treated with CHIR/SB/LY, a notable rise in percentages (Fig. 4A, B) and numbers (Fig. 4C) of phenotypically defined HSCs was observed within the hematopoietic population at day 12 of differentiation compared to controls. To provide corroborative evidence of the positive effect of CHIR/SB/LY on HSPC maturation during iPSC differentiation, we also compared expression of markers recently pinpointed as landmarks of HSC maturity (e.g., PROM1 and HLA-DRA) or immaturity (e.g., CDH5 and MEIS2) in a single cell transcriptomic analysis of human HSPCs at distinct stages of maturation during ontogeny [28]. Addition of CHIR/SB/LY induced robust increased expression of the maturation markers PROM1 and HLA-DRA (Fig. 4D), and concurrent downregulation of the immaturity genes CDH5 and MEIS2 (Fig. 4E) compared to control groups.

To address whether addition of CHIR/SB/LY during iPSC differentiation conferred enhanced functional properties to HSPCs *in vivo*, we quantified early homing (24 hours) and long-term engraftment (4 months) of human cells within hematopoietic tissues of NOD.Cg-Kit^W-41J^ Tyr^+^ Prkdc^scid^ Il2rg^tm1Wjl^/ThomJ (NBSGW) mice transplanted with CD34^+^ HSPCs differentiated from human iPSCS with or without CHIR/SB/LY small molecules. We first measured expression of the chemokine receptor CXCR4 and the β1-integrin VLA-4, two foremost cell surface markers implicated in homing and retention of HSPCs within sites of hematopoiesis after transplantation [52,53]. The proportion of human CD34^+^ HSPCs expressing CXCR4 (Fig. 5A, B) or VLA-4 surface proteins (Fig. 5C, D) significantly increased with addition of CHIR/SB/LY during iPSC differentiation but remained lower than frequencies detected in cultured CD34^+^ HSPCs collected by G-CSF mobilization and apheresis of a healthy subject (Fig. 5A). With CHIR/SB/LY supplementation, expression of both markers was also upregulated, as measured by mean fluorescence intensity on flow cytometry plots (Fig. 5E, F). However, whereas bona fide human CD34^+^ cells were readily detectable within marrow and spleen tissues one day following tail vein injection of NBSGW mice, human cell chimerism was not measurable in mice transplanted with similar numbers of CD34^+^ HSPCs derived from iPSC differentiation cultures (Fig. 5G-J). Consistent with the early homing and retention failure *in vivo*, HSPCs derived from iPSC differentiation cultures in the presence or absence of CHIR/SB/LY did not sustain levels of long-term hematopoietic engraftment comparable to somatic human CD34+ cells after transplantation into NBSGW mouse recipients (Fig. 5K, L and Fig. S2).

## Discussion

In this study, we investigate the long-standing need for optimized human iPSC differentiation systems to enable production of clinical-grade HSCs with functional properties *in vivo* for therapeutic applications. This work stemmed from our previous observation that arterial HE within the non-hematopoietic adherent monolayer that forms during the initial phase of our differentiation platform was scarce and likely insufficient to fully support the generation of bona fide engrafting HSCs in culture [43]. Thus, in keeping with the prevailing arterial-specification model [23–27], we aimed to augment production of arterial HE during iPSC differentiation and evaluated the impact on HSPC formation in culture. By stage-specific addition of small molecules to modulate key signaling pathways independently shown to control arterial fate during vascular development, including WNT, Activin/Nodal and MAPK pathways, we made two observations. First, we found that combined addition of CHIR/SB/LY during human iPSC hematopoietic differentiation could synergistically promote arterial specification of HE, peaking at day 6 of culture, compared to control groups. Second, we showed that this early increase in arterial HE production promoted formation of definitive HSPCs with improved multilineage differentiation capacity, self-renewal potential and maturation state at day 12 of culture. We foresee that adjunct treatment with CHIR/SB/LY during human iPSC differentiation may become routinely used to enhance the arterial HE program and boost definitive multilineage hematopoietic formation in culture.

This work is grounded on the hypothesis that arterial specification is essential for definitive hematopoiesis to arise during ontogeny and in *ex vivo* culture systems. This premise has gained credence based on the observed origin of HSCs within the aortic endothelium during development [20,22,54], the demonstration of shared signaling pathways between arterial fate determination and HSC formation [40,41,55,56], and the selective inhibition of definitive hematopoiesis within the dorsal aorta in a murine knock-out model of the artery-specific Ephrin B2 gene [57]. Yet, an arterial-independent model has also been endorsed whereby hematopoietic progenitors expressing endothelial markers, rather than bona fide endothelial cells, undergo HSC specification after transitioning through the permissive AGM arterial environment of the developing conceptus. This hypothesis is primarily substantiated by observations of HSC formation in animal models engineered to disrupt normal arterial specification *in vivo*. However, a direct measure of HSC long-term engraftment activity was not performed in these studies [41,58–62]. In our work, although enhanced arterial HE formation by CHIR/SB/LY supplementation was insufficient to enable the emergence of third-wave engraftable HSCs in human iPSC culture, these results do not exclude arterial specification as a central prerequisite to initiate the HSC program. Instead, activation of an arterial program in combination with additional cues uncoupled from arterial specification during human PSC differentiation are likely needed for efficient *de novo* generation of human HSCs.

Our study focused on modulating WNT, Nodal/Activin and MAPK intrinsic cellular signaling for hematopoietic development *ex vivo*, but further fine-tuning of these and other pathways will likely be required to overcome the engraftment deficit. For instance, although WNT signaling activation plays an essential role in the induction of mesoderm during development, inhibition of this pathway at later stages of differentiation may also be necessary for HSPC development from hemogenic endothelium [63]. In addition, the impact of MAPK signaling activation on arterial HE specification is primarily mediated through stimulation of NOTCH [23], a critical signaling pathways that requires tight regulation during and following EHT to generate and maintain functional HSCs [25,64,65]. Thus, activation of MAPK signaling by LY in our study, although sufficient to enhance the arterial program in combination with CHIR/SB, may not provide the proper intensity of Notch signal permissive for third-wave HSC specification. Stage-specific modulation of additional intrinsic cellular inputs, such as ectopic expression of single or combinations of transcription factors linked to HSC regulation and inhibition of key epigenetic regulators known to restrict hematopoiesis during development may also be required to promote full functionality of HSCs engineered *ex vivo* [11,66,67]. Importantly, because hematopoietic cells emerge and mature within the three-dimensional niches of specialized anatomical sites during development, integration of extrinsic environmental cues during human PSC differentiation *ex vivo*, namely cell-cell contact, cell-matrix interactions, diffusible factors and biomechanical forces, may further enhance the emergence of definitive hematopoiesis from arterialized HE in culture (reviewed in [11,68]).

Our study also uncovered an early transplantation failure of iPSC-derived HSPCs in xenograft murine models. Although efficient homing, retention and survival within specialized niches of recipient hematopoietic tissues after transplantation are indispensable for long-term engraftment of HSCs, the early homing failure alone is unlikely sufficient to explain the engraftment deficit observed at 4 months in our study. In a recent investigation [69], ectopic expression of the homing receptor CXCR4 improved short-term marrow retention but was insufficient to confer sustained engraftment. The iPSC differentiation platform used in our study enabled production of human HSPCs with CXCR4 and VLA-4 activity independent of ectopic expression, and medium supplementation with CHIR/SB/LY further enhanced expression of both markers. Hence, the inability of these cells to home and survive after transplantation cannot be attributed to limited expression of the CXCR4 chemokine receptor or VLA-4 integrin. A detailed analysis of their functional integrity, including ligand binding affinity, receptor conformational changes, signal transduction, internalization and degradation, is needed to fully understand the mechanism underpinning the inadequate homing capabilities of HSPCs engineered *ex vivo* [70,71]. A comprehensive examination of additional surface molecules implicated in cellular adhesion (e.g., cadherins, selectins, tetraspanins) and their interacting ligands may also impart new leads for the generation of human HSCs with short-term retention capability and long-term engraftment competency *in vivo*.

## Conclusion

Here, we provide evidence that combined addition of CHIR/SB/LY during human iPSC hematopoietic differentiation enhances formation of arterial HE and phenotypically defined definitive HSPCs with self-renewal and multilineage differentiation capacity. Our observation that CHIR/SB/LY treatment alone was insufficient to enable the emergence of third-wave engraftable HSCs from human iPSC culture indicates that further refinement is required. The clinically-relevant and readily adaptable human iPSC differentiation platform presented in this study offers a unique framework to further define and dependably recapitulate *in vitro* the diverse signals driving hematopoiesis *in vivo* to promote formation of bona fide HSCs *in vitro*.

## Acknowledgements

The authors wish to thank: All members of the Larochelle Laboratory for helpful discussions; Dr. David Stroncek and the NIH Department of Transfusion Medicine and Cell Processing Section staff for providing leukapheresis, isolation, and cryopreservation of CD34^+^ cells; the outpatient clinic nursing staff for recruiting normal volunteers and providing administration teaching to healthy subjects; Dr. Julie Holdridge, Dr. James Hawkins and the mouse core facility staff for excellent animal care; Dr. Pradeep Dagur and the NHLBI flow cytometry core facility staff for technical support with sorting; the NHLBI iPSC core facility staff for assistance with iPSC culture. Graphics in Figs. 1A, 2A and 3A were created with BioRender.com.

## Data Availability Statement

All data are incorporated into the article and its online supplementary material.

