## Supplementary file for "Modulation of WNT, Activin/Nodal and MAPK Signaling Pathways Increases Arterial Hemogenic Endothelium and Hematopoietic Stem/Progenitor Cell Formation During Human iPSC Differentiation"

(1) Cellular and Molecular Therapeutics Branch, National Heart, Lung and Blood Institute (NHLBI), National Institutes of Health (NIH), Bethesda, MD 20892, USA.
(2) iPSC Core Facility, NHLBI, NIH, Bethesda, MD 20892, USA.

*** Corresponding author:**

Andre Larochelle, M.D. Ph.D., National Heart, Lung and Blood Institute, National Institutes of Health, Bethesda, 9000 Rockville, Bethesda, MD 20892, USA.

☎ (301) 814-8289

**Running Title:** Hematopoietic Differentiation of Human iPSCs

**Authors' contribution:**

- **Author 1 (Y.L.):** Conception and design, collection and assembly of data, data analysis and interpretation, manuscript writing, final approval of manuscript.
- **Author 2 (J.D.):** Collection and assembly of data, final approval of manuscript.
- **Author 3 (D.A.):** Collection and assembly of data, final approval of manuscript.
- **Author 4 (J.Z.):** Collection and assembly of data, final approval of manuscript.
- **Author 5 (A.L.):** Conception and design, data analysis and interpretation, manuscript writing, final approval of manuscript.

**Funding:**

This work was supported by the Intramural Research Program of the National Heart, Lung, and Blood Institute, National Institutes of Health (Z99 HL999999 and ZIA HL006217).

**Keywords:**

Induced pluripotent stem cells, Hematopoietic stem/progenitor cells, Arterial hemogenic endothelium, Activin/Nodal, WNT and MAPK signaling pathways.

This file includes:

- **Supplementary Figures**

  - **Figure S1**: Modulation of WNT, Activin/Nodal, and MAPK signaling pathways during human iPSC differentiation enhances formation of arterial hemogenic endothelium.
  - **Figure S2**: Optimization of human iPSC differentiation for maximal production of arterial hemogenic endothelium via modulation of WNT, Activin/Nodal, and MAPK signaling pathways.
  - **Figure S3:** Modulation of WNT, Activin/Nodal, and MAPK signaling pathways during human iPSC differentiation enhances formation of hematopoietic progenitors.
  - **Figure S4:** Modulation of WNT, Activin/Nodal, and MAPK signaling pathways during human iPSC differentiation enhances formation of HSPCs with self-renewal capacity.
  - **Figure S5:** Modulation of WNT, Activin/Nodal, and MAPK signaling pathways during human iPSC differentiation enhances formation of HSPCs with phenotypic attributes of maturation.
  - **Figure S6:** Long-term *in vivo* engraftment potential of human iPSC-derived CD34+ HSPCs.

- **Supplementary Tables**

  - **Table S1**: Fluorochrome-conjugated monoclonal antibodies for flow cytometry.
  - **Table S2**: Primers for real-time quantitative PCR.

#### Supplementary Figures

##### Figure S1

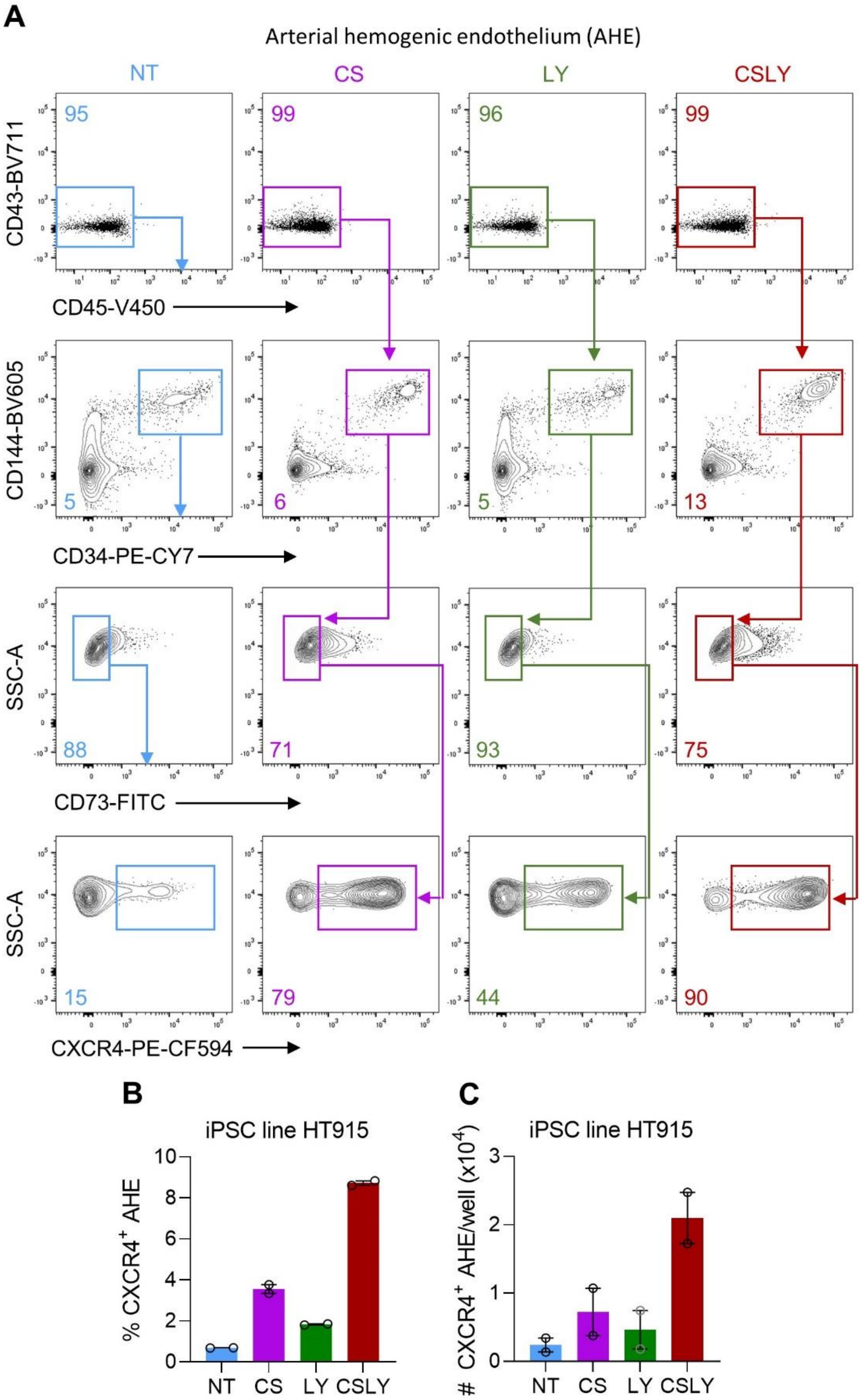

**Figure S1 | Modulation of WNT, Activin/Nodal, and MAPK signaling pathways during human iPSC differentiation enhances formation of arterial hemogenic endothelium.** Human iPSCs were differentiated as illustrated in Fig. 1A. All analyses were performed with cells collected at day 6 of differentiation in the absence of small molecule adjunct treatment (non-treated, NT) or in the presence of CHIR/SB (CS), LY or CHIR/SB/LY (CSLY). **(A)** Representative flow cytometry plots of arterial hemogenic endothelium (AHE)-defining markers ($CD43^-CD45^-CD144^+CD34^+CD73^-CXCR4^+$ or $DLL4^+$). Data shown in this figure were obtained with human iPSC line HT914. **(B)** Percentages of $CXCR4^+$ AHE cells. (**C)** Absolute numbers of $CXCR4^+$ AHE cells per culture well. In panels B and C, data are displayed as mean ± standard error of the mean (SEM) of 2 independent experiments. Associated with Fig.1.

**Figure S2**

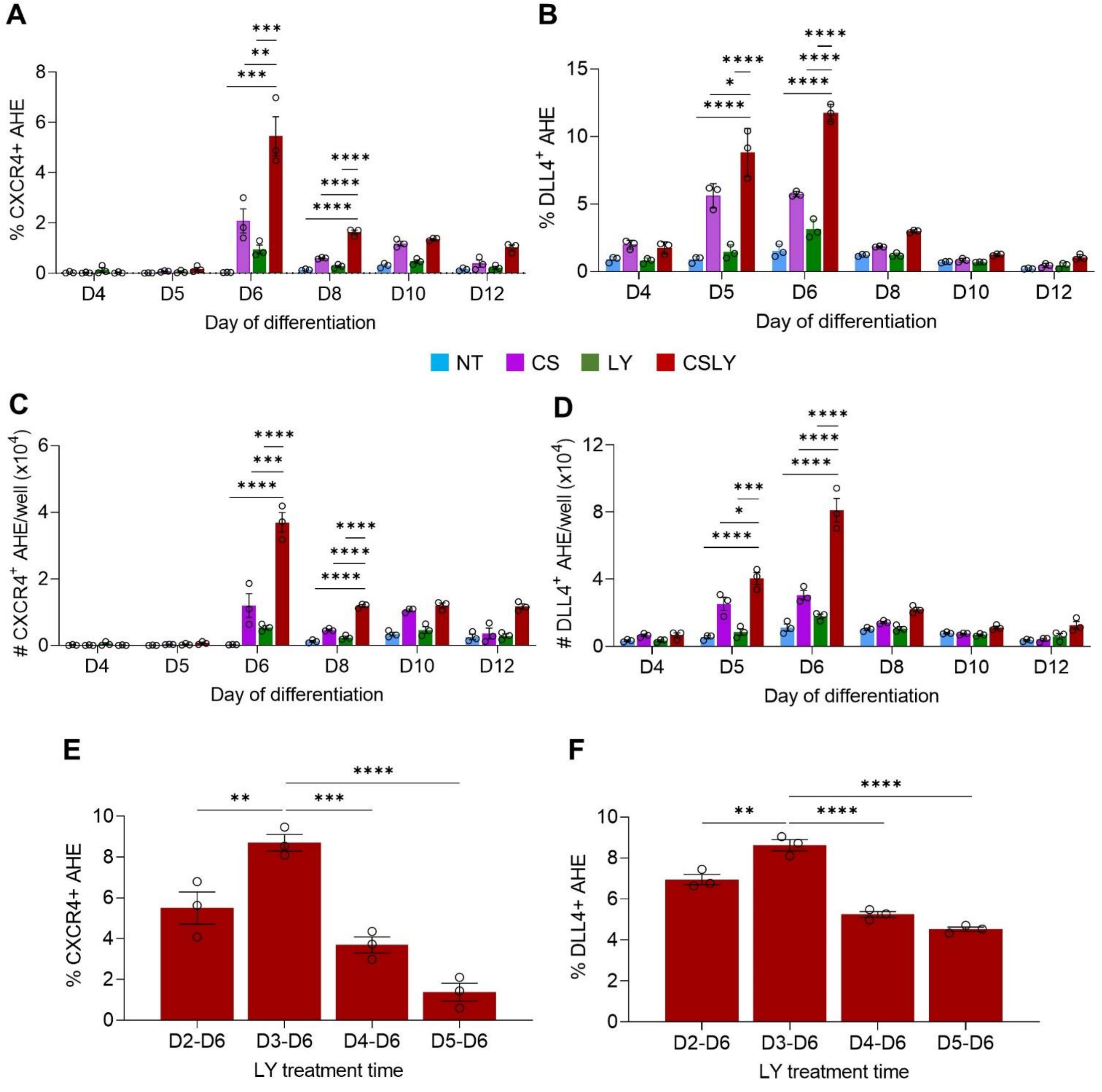
A
% CXCR4+ AHE
Day of differentiation
D4
D5
D6
D8
D10
D12
B
% DLL4+ AHE
Day of differentiation
NT
CS
LY
CSLY
C
### CXCR4+ AHE/well (x10^4)
D
### DLL4+ AHE/well (x10^4)
E
% CXCR4+ AHE
LY treatment time
D2-D6
D3-D6
D4-D6
D5-D6
F
% DLL4+ AHE
LY treatment time

**Figure S2 | Optimization of human iPSC differentiation for maximal production of arterial hemogenic endothelium via modulation of WNT, Activin/Nodal, and MAPK signaling pathways.** Human iPSCs were differentiated as illustrated in Fig. 1A in the absence of small molecule adjunct treatment (non-treated, NT) or in the presence of CHIR/SB (CS), LY or CHIR/SB/LY (CSLY). **(A-D)** Determination of the peak effect of CSLY addition on arterial hemogenic endothelium (AHE) production during human iPSC differentiation. Maximal AHE formation was observed at day 6 of differentiation. Panels display percentages of $CXCR4^+$ AHE cells (A), percentages of $DLL4^+$ AHE cells (B), absolute numbers of $CXCR4^+$ AHE cells per culture well (C), and absolute numbers of $DLL4^+$ AHE cells per culture well. **(E-F)** Determination of the optimal LY treatment scheme for maximal AHE production during human iPSC differentiation. All analyses were performed with cells collected at day 6 of differentiation with indicated treatments. Maximal AHE formation was observed when LY was supplemented from day 3 through day 6 of culture. Panels display percentages of $CXCR4^+$ AHE cells (E), and percentages of $DLL4^+$ AHE cells (F). Data shown in this figure were obtained with human iPSC line MCND-TEN-S2. Data are displayed as mean ± SEM of 3 independent experiments. One-way ordinary ANOVA test with Dunnett correction was used. * $p \leq 0.05$, ** $p \leq 0.01$, *** $p \leq 0.001$, **** $p \leq 0.0001$. Associated with Fig.1.

**Figure S3**

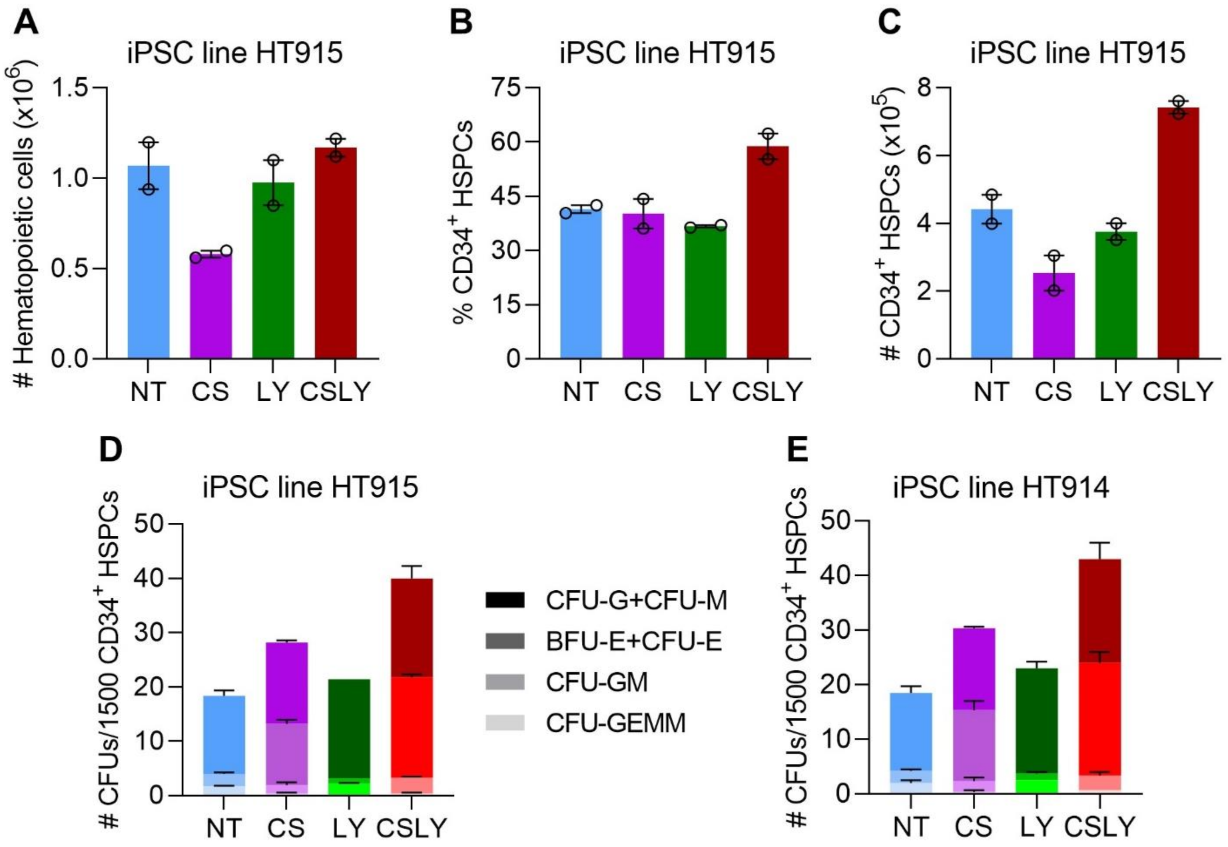
A
iPSC line HT915
### Hematopoietic cells (x10^6)
1.5
1.0
0.5
0.0
NT
CS
LY
CSLY
B
iPSC line HT915
% CD34+ HSPCs
75
60
45
30
15
0
NT
CS
LY
CSLY
C
iPSC line HT915
### CD34+ HSPCs (x10^5)
8
6
4
2
0
NT
CS
LY
CSLY
D
iPSC line HT915
### CFUs/1500 CD34+ HSPCs
50
40
30
20
10
0
NT
CS
LY
CSLY
CFU-G+CFU-M
BFU-E+CFU-E
CFU-GM
CFU-GEMM
E
iPSC line HT914
### CFUs/1500 CD34+ HSPCs
50
40
30
20
10
0
NT
CS
LY
CSLY

**Figure S3 | Modulation of WNT, Activin/Nodal, and MAPK signaling pathways during human iPSC differentiation enhances formation of hematopoietic progenitors.** Human iPSCs were differentiated as illustrated in Fig. 1A in the absence of small molecule adjunct treatment (non-treated, NT) or in the presence of CHIR/SB (CS), LY or CHIR/SB/LY (CSLY). Hematopoietic progenitor activity of cells released within the culture supernatant at day 12 of differentiation with indicated treatments was evaluated by flow cytometry and colony forming unit (CFU) assay. (**A)** Absolute numbers of $CD43^{+}CD45^{+/-}$ hematopoietic cells per culture well. **(B)** Percentages of $CD34^{+}$ HSPCs. **(C)** Absolute numbers of $CD34^{+}$ HSPCs per culture well. **(D, E)** Numbers of myeloid (CFU-G, CFU-M, CFU-GM and CFU-GEMM) and erythroid (CFU-E, BFU-E) colonies per 1500 CD34+ cells purified by FACS. Data shown in this figure were obtained with human iPSC lines HT915 (panels A-D) and HT914 (panel E). Data are displayed as mean ± SEM of 2 independent experiments. Associated with Fig. 2.

**Figure S4**

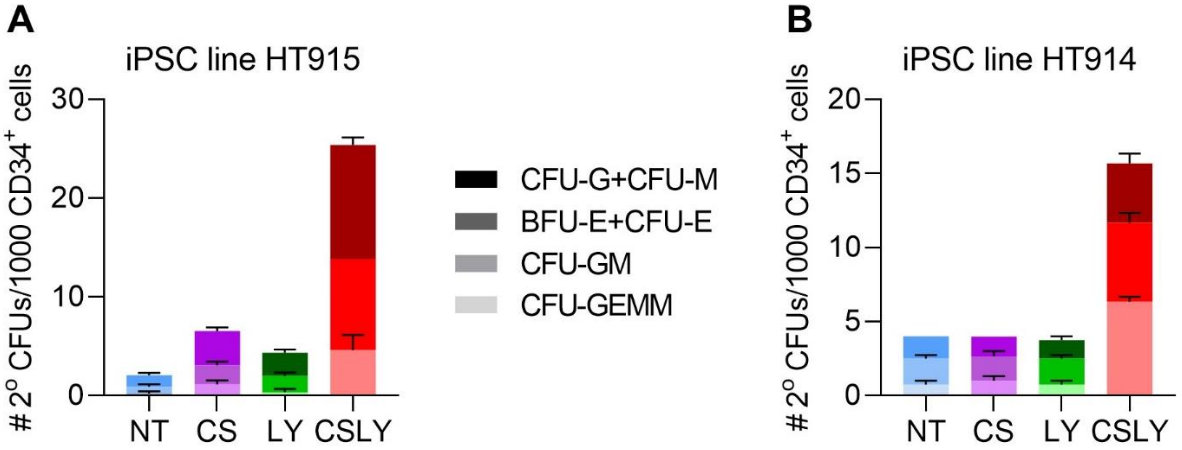
A
iPSC line HT915
### 2$^{o}$ CFUs/1000 CD34$^{+}$ cells
30
20
10
0
NT
CS
LY
CSLY
CFU-G+CFU-M
BFU-E+CFU-E
CFU-GM
CFU-GEMM
B
iPSC line HT914
### 2$^{o}$ CFUs/1000 CD34$^{+}$ cells
20
15
10
5
0
NT
CS
LY
CSLY

**Figure S4 | Modulation of WNT, Activin/Nodal, and MAPK signaling pathways during human iPSC differentiation enhances formation of HSPCs with self-renewal capacity.** Human iPSCs were differentiated as illustrated in Fig. 1A in the absence of small molecule adjunct treatment (non-treated, NT) or in the presence of CHIR/SB (CS), LY or CHIR/SB/LY (CSLY). Self-renewal capacity of HSPCs released within the culture supernatant was evaluated at day 12 of differentiation with indicated treatments. (**A, B)** Secondary clonogenic assay. A total of 1000 CD34+ HSPCs purified by FACS were plated in primary CFU assays. After 12-14 days, colonies were scored, pooled and equal numbers of cells were replated for each condition. Secondary CFU plates were scored at day 12-14 for myeloid (CFU-G, CFU-M, CFU-GM and CFU-GEMM) and erythroid (CFU-E, BFU-E) colonies, and counts were normalized to the total number of cells from primary CFU plates. Data shown in this figure were obtained with human iPSC lines HT915 (panel A) and HT914 (panel B). Data are displayed as mean ± SEM of 2 independent experiments. Associated with Fig. 3.

**Figure S5**

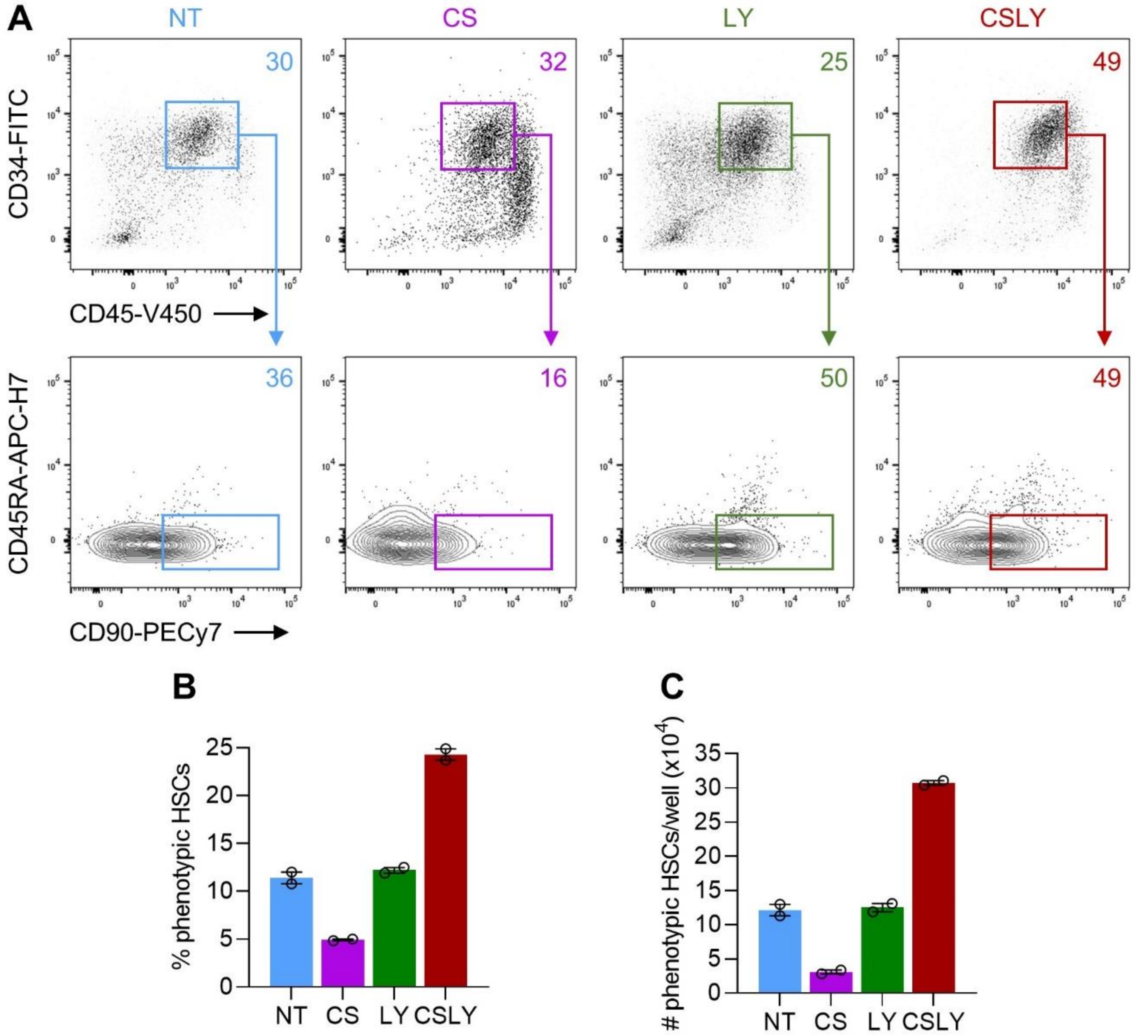
A
NT
CS
LY
CSLY
30
32
25
49
CD34-FITC
CD45-V450
36
16
50
49
CD45RA-APC-H7
CD90-PECy7
B
% phenotypic HSCs
NT CS LY CSLY
C
### phenotypic HSCs/well (x10^4)
NT CS LY CSLY

**Figure S5 | Modulation of WNT, Activin/Nodal, and MAPK signaling pathways during human iPSC differentiation enhances formation of HSPCs with phenotypic attributes of maturation.** Human iPSCs were differentiated as illustrated in Fig. 1A in the absence of small molecule adjunct treatment (non-treated, NT) or in the presence of CHIR/SB (CS), LY or CHIR/SB/LY (CSLY). All analyses were performed with HSPCs collected at day 12 of differentiation with indicated treatments. **(A)** Representative flow cytometry plots of phenotypically defined HSCs ($CD45^{+}CD34^{+}CD45RA^{-}CD90^{+}$). **(B)** Percentages of phenotypically defined HSCs. **(C)** Absolute numbers of phenotypically defined HSCs per culture well. Data shown in this figure were obtained with human iPSC line HT915. In panels B and C, data are displayed as mean ± SEM of 2 independent experiments. Associated with Fig. 4.

**Figure S6**

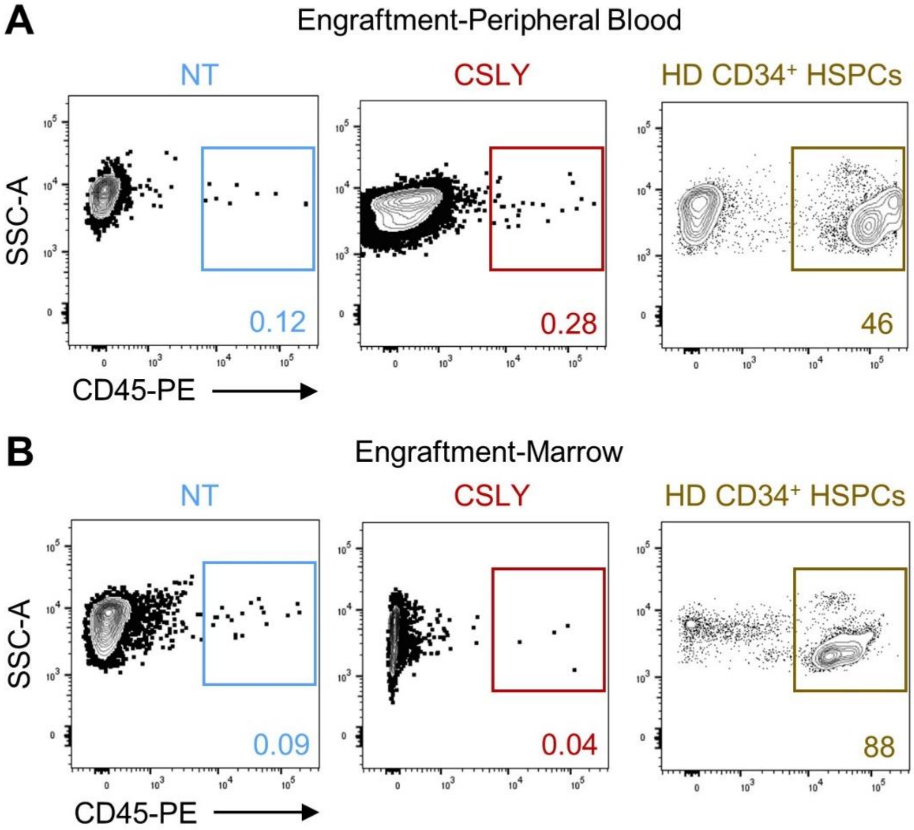

A
Engraftment-Peripheral Blood
NT
CSLY
HD CD34⁺ HSPCs
SSC-A
0.12
0.28
46
CD45-PE
B
Engraftment-Marrow
NT
CSLY
HD CD34⁺ HSPCs
SSC-A
0.09
0.04
88
CD45-PE

**Figure S6 | Long-term *in vivo* engraftment potential of human iPSC-derived CD34+ HSPCs.** Human iPSCs were differentiated as illustrated in Fig. 1A in the absence of small molecule adjunct treatment (non-treated, NT) or in the presence of CHIR/SB/LY (CSLY). All analyses were performed using $CD34^+$ HSPCs purified by FACS from the culture supernatant at day 12 of differentiation with indicated treatments or collected by G-CSF mobilization and apheresis of a healthy donor (HD). **(A)** Representative flow cytometry plots of human $CD45^+$ cells detected within the peripheral blood of NBSGW mice at 12 weeks post-transplantation. **(B)** Representative flow cytometry plots of human $CD45^+$ cells detected within the marrow of NBSGW mice at 16 weeks post-transplantation. Data shown in this figure were obtained with human iPSC line MCND-TEN-S2. Associated with Fig. 5.

#### Supplementary Tables

**Table S1 | Fluorochrome-conjugated monoclonal antibodies for flow cytometry**

| Antibody Conjugate | Clone | Supplier | Catalog number | Species |
|---|---|---|---|---|
| CD34-PE-Cy7 | 581 | BD Pharmingen | 560710 | Mouse anti-human |
| CD34-FITC | 581 | BD Bioscience | 555821 | Mouse anti-human |
| CD34-APC | 8G12 | BD Bioscience | 345804 | Mouse anti-human |
| CD43-BV711 | 1G10 | BD OptiBuild™ | 743614 | Mouse anti-human |
| CD45-V450 | HI30 | BD Horizon | 560367 | Mouse anti-human |
| CD45-PE | HI30 | BD Bioscience | 555483 | Mouse anti-human |
| CD45RA-APC-H7 | HI100 | BD Bioscience | 560674 | Mouse anti-human |
| CD73-FITC | AD2 | BD Pharmingen | 561254 | Mouse anti-human |
| CD90-PECy7 | 5E10 | BD Pharmingen | 561558 | Mouse anti-human |
| CD144-BV605 | 55-7H1 | BD OptiBuild™ | 743705 | Mouse anti-human |
| CXCR4-PE-CF594 | 12G5 | BD Horizon | 562389 | Mouse anti-human |
| DLL4-APC | MOPC-21 | Biolegend | 346507 | Mouse anti-human |
| VLA-4-BV421 | 9F10 | BD Bioscience | 565277 | Mouse anti-human |

**Table S2 | Primers for real-time quantitative PCR**

| Primer Name | Primer Sequence |
|---|---|
| PROM1-F | 5’ AGTCGGAAACTGGCAGATAGC 3’ |
| PROM1-R | 5’ GGTAGTGTTGTACTGGGCCAAT 3’ |
| HLA-DRA-F | 5’ ATGGCCATAAGTGGAGTCCC 3’ |
| HLA-DRA-R | 5’ CTCCATGTGCCTTACAGAGG 3’ |
| CDH5-F | 5’ AAACACCTCACTTCCCCATC 3’ |
| CDH5-R | 5’ ACCTTGCCCACATATTCTCC 3’ |
| MEIS2-F | 5’ TCCACAAATCTCGCTGACC 3’ |
| MEIS2-R | 5’ GTCTAAACCATCCCCTTGCTC 3’ |
| GAPDH-F | 5’ ACATCGCTCAGACACCATG 3’ |
| GAPDH-R | 5’ TGTAGTTGAGGTCAATGAAGGG 3’ |

F: Forward primer; R: Reverse primer
